# Magnetic particle imaging of magnetotactic bacteria as living contrast agents is improved by altering magnetosome structures

**DOI:** 10.1101/2021.12.03.471101

**Authors:** Ashley V. Makela, Melissa A. Schott, Cody Madsen, Emily Greeson, Christopher H. Contag

**Affiliations:** Institute for Quantitative Health Science and Engineering, Michigan State University, East Lansing, MI, USA; Homer Stryker M.D. School of Medicine, Western Michigan University, Kalamazoo, MI, USA; Department of Biomedical Engineering, Michigan State University, East Lansing, MI, USA; Department of Microbiology and Molecular Genetics, Michigan State University, East Lansing, MI, USA

**Keywords:** magnetotactic bacteria, magnetic particle imaging, magnetosomes, bioluminescence, iron nanoparticle

## Abstract

Iron nanoparticles used as imaging contrast agents can help differentiate between normal and diseased tissue, or track cell movement and localize pathologies. Magnetic particle imaging (MPI) is an imaging modality that uses the magnetic properties of iron nanoparticles to provide specific, quantitative and sensitive imaging data. MPI signals depend on the size, structure and composition of the nanoparticles; MPI-tailored nanoparticles have been developed by modifying these properties. Magnetotactic bacteria produce magnetosomes which mimic synthetic nanoparticles, and thus comprise a living contrast agent in which nanoparticle formation can be modified by mutating genes. Specifically, genes that encode proteins critical to magnetosome formation and regulation, such as mamJ which helps with filament turnover. Deletion of mamJ in Magnetospirillum gryphiswaldense, MSR-1 led to clustered magnetosomes instead of the typical linear chains. Here we examined the effects of this magnetosome structure and revealed improved MPI signal and resolution from clustered magnetosomes compared to linear chains. Bioluminescent MSR-1 with the mamJ deletion were injected intravenously into tumor-bearing and healthy mice and imaged using both in vivo bioluminescence imaging (BLI) and MPI. BLI revealed the location and viability of bacteria which was used to validate localization of MPI signals. BLI identified the viability of MSR-1 for 24 hours and MPI detected iron in the liver and in multiple tumors. Development of living contrast agents offers new opportunities for imaging and therapy by using multimodality imaging to track the location and viability of the therapy and the resulting biological effects.

---

Contrast agents have distinctive molecular or structural signatures that are used in biomedical imaging and composition, shape and size can be used to enhance these distinctive signatures. Such agents are often required in imaging studies to differentiate between normal and diseased tissue, to track cell movement or to localize pathologies^1^. Magnetic nanoparticles (MNP) have commonly been used as imaging contrast agents in applications of magnetic resonance imaging (MRI)^2,3^ and more recently, magnetic particle imaging (MPI)^4–10^. MRI visualizes the MNP as a signal void, due to its magnetic field inducing a local change in the relaxation of surrounding protons. MPI is an emerging imaging technique which detects iron particles at concentrations approaching a picogram without signal attenuation as depth in the tissue increases. MPI images are created using a gradient magnetic field consisting of a field free region (FFR) and an alternating magnetic field (AMF). The FFR rasters through a prescribed region of interest, and as the FFR moves across the MNP there is a change in magnetization, inducing a voltage received by the system which is then transformed into signal^11–13^. MPI allows for quantification because the signal is linearly proportional to the amount of iron present. If there is prior knowledge of the amount of iron per cell, this information can further provide a means of calculating the number of iron-containing cells at a given anatomic site.

The use of synthetic MNPs to label cells is limited by repeated dilution of the MNP associated with cell division. Each progeny cell will contain less iron after cell division, resulting in reduced signal which will eventually fall below imaging detection thresholds^14^. The use of a replicating, biologically produced iron nanoparticle could address this limitation. Magnetotactic bacteria are organisms with the unique ability to biomineralize iron into cuboctahedral magnetite (Fe_3_O_4_) crystals compartmentalized intracellularly in magnetosomes^15,16^. In nature, the magnetosomes allow these bacteria to perform magnetotaxis^17^. Magnetotactic bacteria maintain their magnetic signatures through continued synthesis of magnetosomes from intracellular iron under low oxygen conditions^18^. Magnetotactic bacteria have been explored in the fields of imaging and bacteriotherapy for magnetic drug-targeted therapies, hyperthermia treatments, and other non-therapeutic applications such as biosensors^18–20^. Here we expand on the use of magnetotactic bacteria in the field of molecular imaging and investigate the magnetic magnetosomes in the development of biological contrast agents for MPI.

Development of MNP specifically for MPI applications include MNP that form aggregates or clusters which are used in MPI^21^ and AMF^22^ applications. Magnetosome synthesis in magnetotactic bacteria has been related to at least 100 genes, which are organized into five operons which make up the magnetosome island of the genome^23,24^. Genetic deletions or knockouts within the magnetosome island have been shown to affect magnetosome assembly and structure^25,26^. Deletions of the actin-like cytoskeletal filament *mamK* and filament turnover promoting *mamJ* genes, which are responsible for magnetosome chain assembly, have shown to affect the overall magnetosome structures^27–29^. Here, we examine the effects of a *mamJ* knockout (Δ*mamJ*) and the resulting magnetosome configuration on MPI signals. The *mamJ* gene encodes an acidic protein that is involved in formation and maintenance of a chain-like structure of magnetosomes that is observed in *Magnetospirillum spp*.^27,28,30^. The *mamJ* knockout changes the structure of the magnetosome chain into a cluster^27^ closely resembling a multicore, synthetic MNP^31^.

Since magnetotactic bacteria are living contrast agents, an orthogonal measure of their viability would be useful for monitoring their development as MPI contrast agents. Luciferases are metabolic reporters, dependent on oxygen and energy for their activity, and are the foundation of *in vivo* bioluminescence imaging (BLI)^32^. The use of BLI in conjunction with MPI would enable following the viability of magnetotactic bacteria *in vivo* and correlating these data with the MPI images. The engineered *luxA-E* bacterial operon is based on the native *luxCDABE* operon^33^ and encodes five enzymes for bioluminescence. By introducing the *luxA-E* operon into *M. gryphiswaldense* MSR-1, cellular viability can be monitored using BLI without the need of an added substrate.

A living bacterial contrast agent could be applied as a bacteriotherapeutic. Bacteriotherapies have become relevant in cancer treatment, such as *Bacille Calmette-Guerin* being used to effectively treat bladder carcinoma^34,35^ through what is assumed to be an activation of the immune response. Further, genetically modified *Listeria monocytogenes* is in Phase 1 and Phase 2 clinical trials as a bacteria-based vaccine to treat various types of cancer^36,37^. Likewise, magnetotactic bacteria have also been investigated for use in bacteriotherapy^38^. The effective use of these bacteriotherapies and their documented safety provide assurance when looking towards the future of applying magnetotactic bacteria in this context. Magnetotactic bacteria could be used as heating agents to eliminate cells, using magnetohyperthermia^39,40^, or could be engineered and induced to express therapeutic genes^41^, targeting the tumor microenvironment to elicit anti-tumoral properties.

## Results and discussion

### Phenotypic differences in MSR-1 strains

The two strains of MSR-1 were morphologically different; wild type (WT) contained long chains of magnetosomes (**Figure 1A**) that usually spanned the entire length of the bacterium while Δ*mamJ* contained one or two dense clusters of magnetosomes^27^ (**Figure 1B**). The Δ*mamJ* strain had significantly less iron per cell than the WT strain (**Figure 1C**) with 8.17 +/- 1.24 and 13.22 +/- 0.25 fg/cell, respectively (*p*=.0023) as determined using Inductively Coupled Plasma-Mass Spectrometry (ICP-MS), under the same growth conditions.

**Figure 1.**
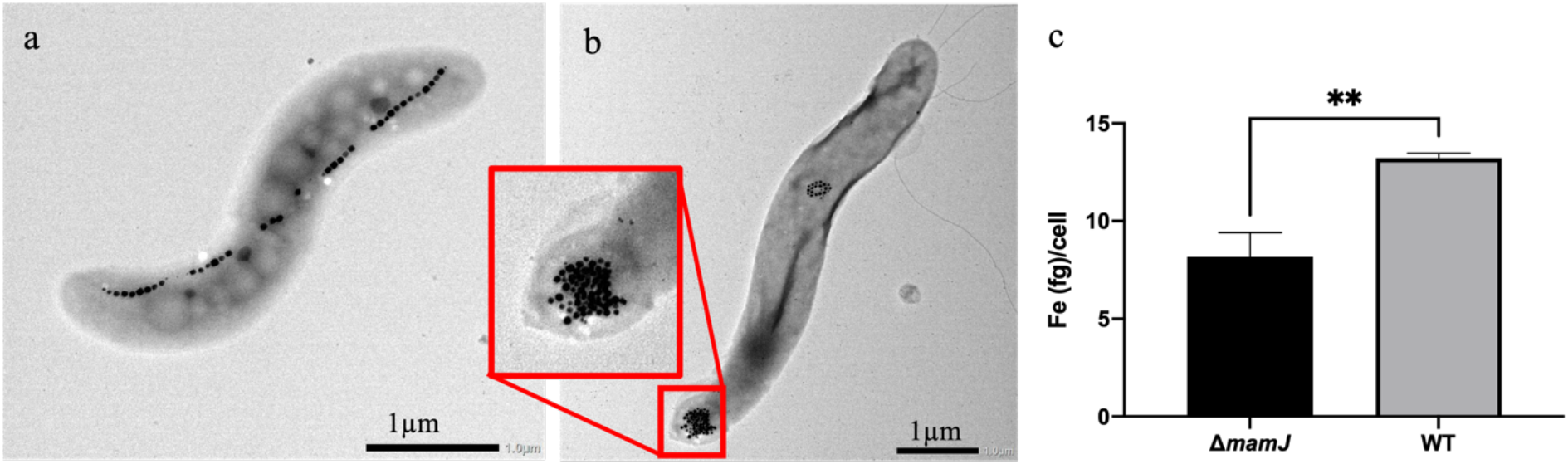
Phenotypic and iron content differences in Magnetospirillum gryphiswaldense MSR-1 WT and *ΔmamJ*. MSR-1 WT (a) and *ΔmamJ* (b) contain chains and clusters of magnetosomes, respectively. ICP-MS analysis was used to calculate iron (Fe) per cell, identifying significantly more Fe/cell in the WT compared to *ΔmamJ* (c, n = 3). **p<.01

### Altered magnetosomes change magnetic particle imaging signal

The same number of either WT or Δ*mamJ* cells were imaged by MPI, as pellets in phosphate buffered saline (PBS), and differences in signal intensity and resolution were identified. The Δ*mamJ* strain (**Figure 2A**) shows clear, defined signal localized to the pellet. Conversely, the WT strain (**Figure 2B**) shows a decrease in signal at the site of the pellet with some signal extending along the y-axis. When normalized to iron content, as determined by ICP-MS, there was significantly more (*p*=.0012) MPI signal generated from the Δ*mamJ* strain (221.2 +/- 26.19 a.u./*μ*g Fe) vs the WT strain (58.79 +/- 22.25 a.u./*μ*g Fe). MPI relaxometry was then used to determine spatial resolution of each strain’s magnetic particles. Figure 2D compares the point spread function (PSF) of the Δ*mamJ* and WT strains. The resolution of the Δ*mamJ* strain is 2.98 mm whereas the resolution of the WT strain is 7.54 mm.

**Figure 2.**
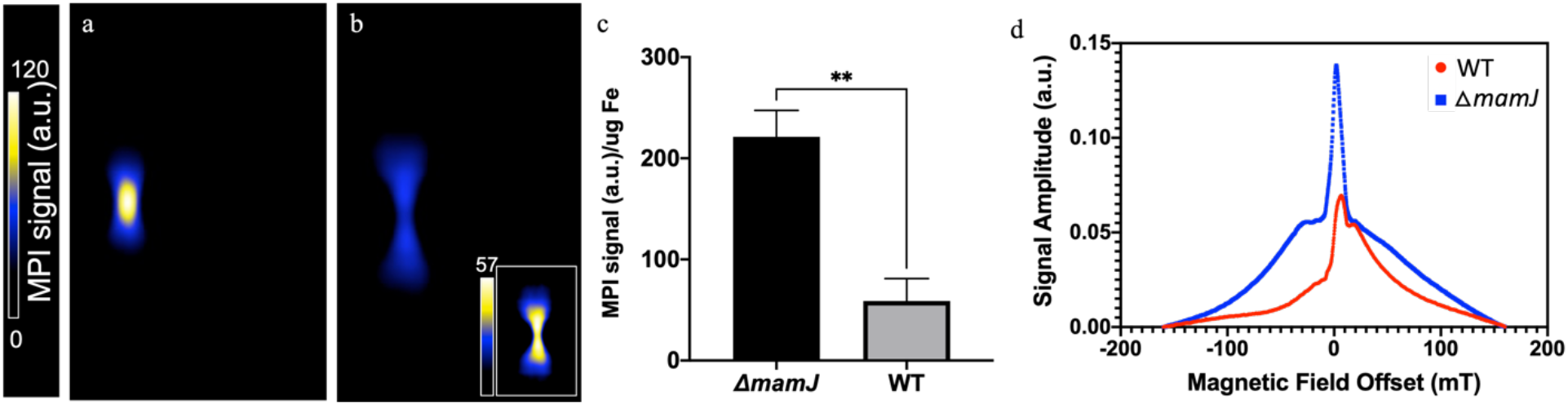
Magnetic Particle Imaging of MSR-1 WT and *ΔmamJ* strains. MPI of *ΔmamJ* (a) and WT (b) pellets. MPI signal normalized to iron content demonstrates an increase in *ΔmamJ* signal compared to WT (c, n = 3). Relaxometry of each strain demonstrates the point spread function created by either *ΔmamJ* or WT, allowing for a measure of signal strength and resolution (d). **p<.01.

Altering the configuration of synthetic iron nanoparticles has resulted in significant changes in the signal strength and resolution when imaged using MPI^44,45^. Here we extend these studies by showing that affect can be achieved using biological iron nanoparticles. Genetically modifying MSR-1 to change the magnetosome configuration, from a linear to a clustered organization^27^, resulted in improved MPI signal strength and resolution. Since magnetosome formation is modulated by proteins encoded by approximately one hundred genes in the magnetosome island in the *M. gryphiswaldense* genome^23,24^, further improvements in MPI signals could be obtained by altering characteristics of the magnetosomes such as the size of individual magnetosomes and additional changes to their structure. Such improvements could enable imaging and detection of magnetotactic bacteria, and this could lead to new *in vivo* applications of this biological tracer.

### Correlation of MPI and bioluminescence signal to bacteria number

MPI signal and cell number were correlated to bioluminescence for each strain. When bacteria were resuspended in media, MPI signal was identified in all numbers of Δ*mamJ luxA-E* (2.5, 1 and 0.5×10^8^) but not WT. The number of bacteria is correlated to bioluminescence for both WT and Δ*mamJ luxA-E* (R^2^ = 0.991 and R^2^ = 0.996, respectively; **Figure 3b**). However, bioluminescence observed from the WT was significantly higher than that of Δ*mamJ luxA-E*, for the same number of cells (*p*<.0001). A linear correlation between MPI signal and bioluminescence was also observed (R^2^ = 0.907) in the Δ*mamJ luxA-E* strain (**Figure 3c**). This could not be calculated in the WT strain as no MPI signal was detected.

**Figure 3.**
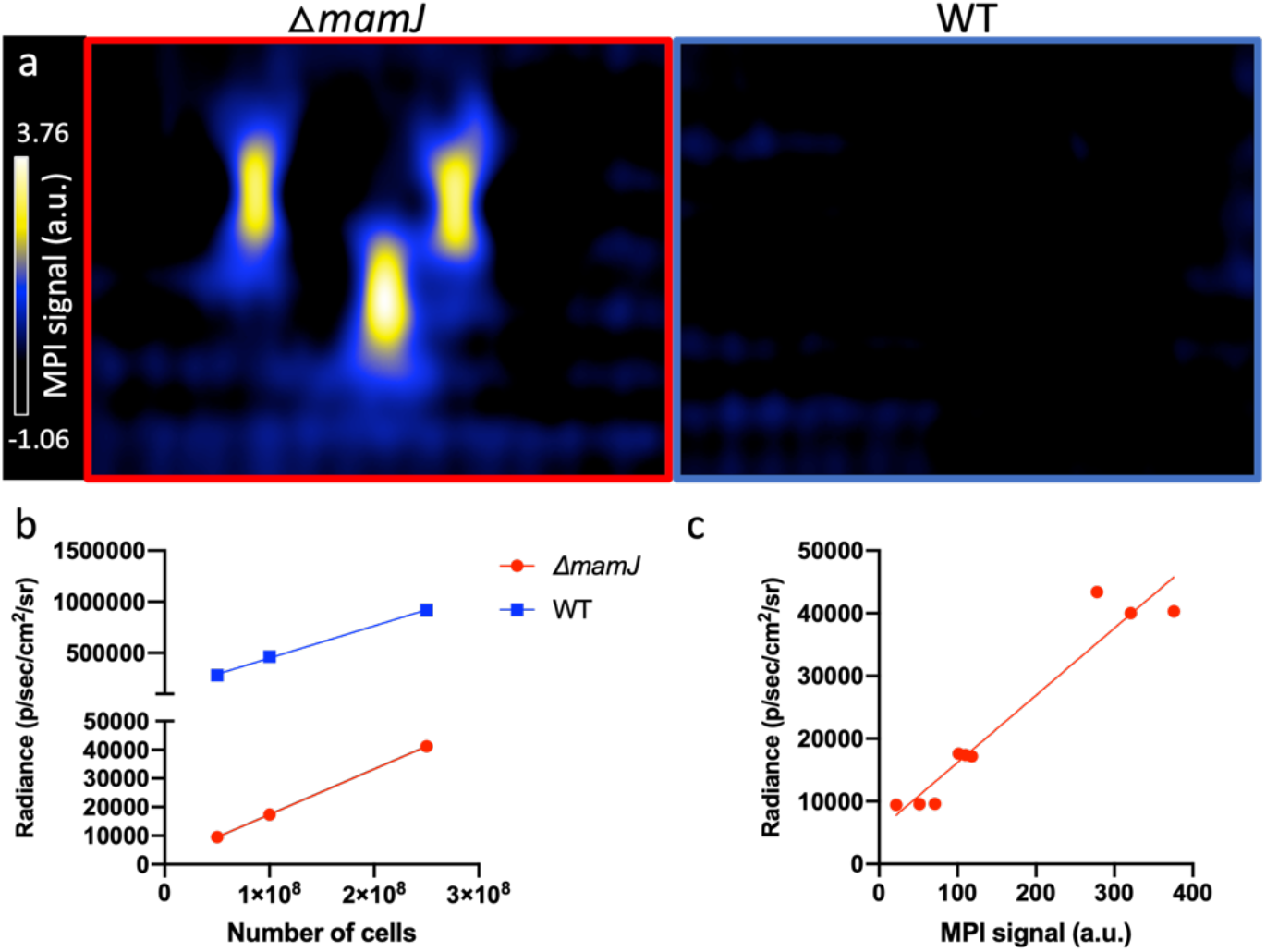
Correlation between bacteria number, luminescence and MPI signal. At the highest cell number only MSR-1 *ΔmamJ luxA-E* produced MPI signal (a). Luminescence was linearly correlated to MPI signal for both *ΔmamJ luxA-E* (R^2^ = 0.9956, n = 3) and WT (R^2^ = 0.991, n = 3) strains (b). WT luminescence was significantly higher than *ΔmamJ luxA-E* for all numbers of cells (p<.0001). *ΔmamJ luxA-E* luminescence is correlated to MPI signal (R^2^ = 0.9295) produced from the same sample (c).

### *In vivo* MPI and BLI of MSR-1 as a living contrast agent

Δ*mamJ luxA-E* were injected intravenously (IV) into healthy mice. BLI and MPI were performed at 5 minutes (m), 1 hour (h) and 24 h post-injection (**Figure 4**) and signals were normalized to the average MPI signal from the Δ*mamJ luxA-E* pellets prior to injection. Five minutes post-injection, bioluminescent signals were apparent throughout the mouse and MPI signals began to appear in the liver. The percent injected dose, as determined by comparing MPI signal with that identified in the pellets prior to injection, were 42.79 +/- 28.09%, 30.34 +/- 1.03% and 62.71 +/- 20.67% at 5m, 1 h and 24 h, respectively. At 1 h post-injection, bioluminescence signals visibly decreased and at 24 h, there was no detectable bioluminescent signals from the mouse.

**Figure 4.**
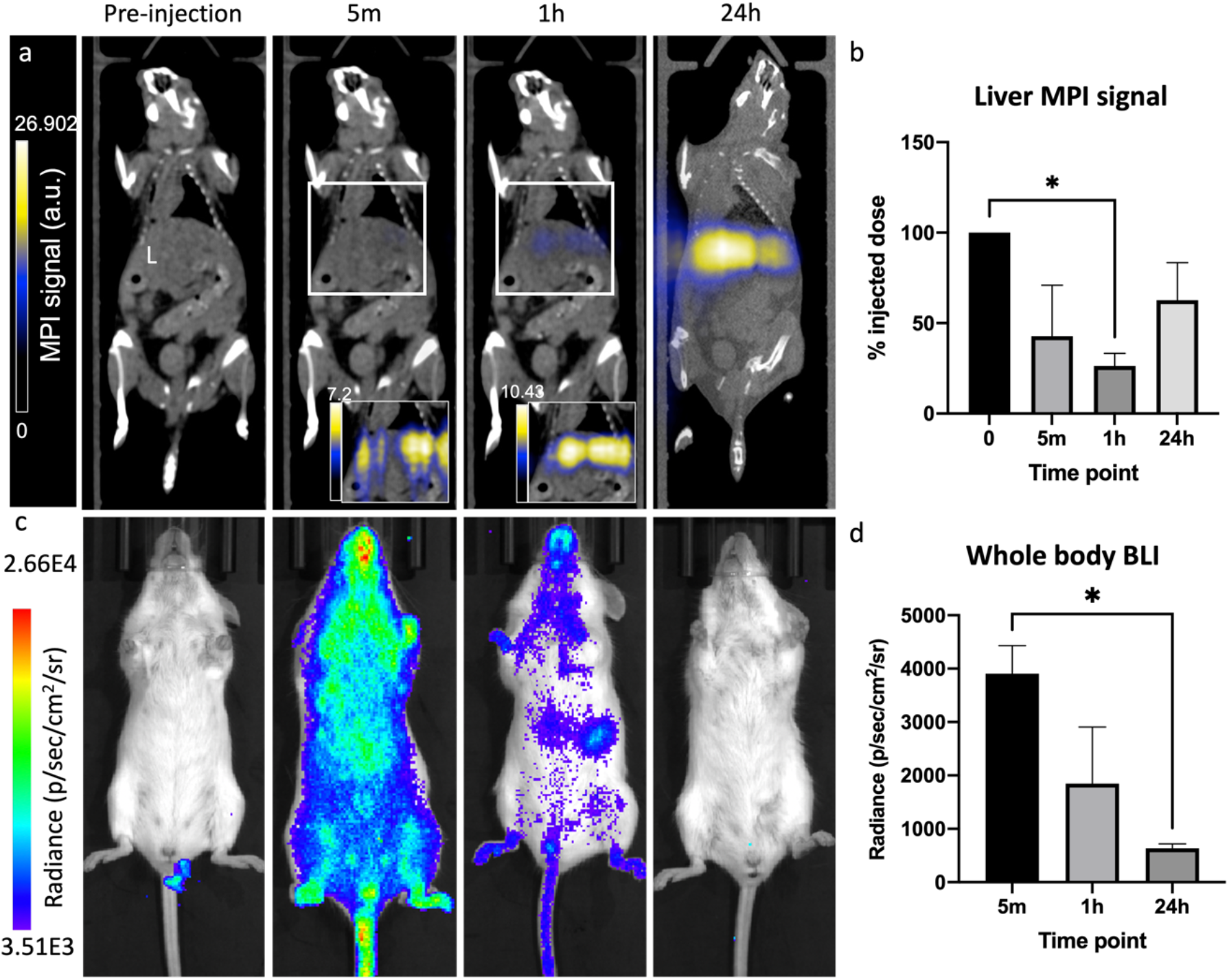
Biodistribution of MSR-1 *ΔmamJ luxA-E* injected intravenously into healthy mice. MPI and BLI signals were monitored at 5 m, 1 h and 24 h post-intravenous injection. MPI signal accumulated in the liver (L) over time, with no signal identified anywhere else (a). MPI signal was used to quantify iron content in the liver at each time point (b). BLI signal was visualized throughout the body at 5 m post-injection, with signal decreasing at 1 h and no signals were detectable at 24 h (c,d). *p<.05

Imaging of excised organs confirmed that the majority of iron aggregated in the liver and some in the spleen (**Figure 5a**). In tumor-bearing mice, a similar *in vivo* distribution was seen after IV injection of Δ*mamJ luxA-E*. The percent injected dose, identified by MPI quantification, which accumulated in the liver was 65.63 +/- 11.14%, 69.11 +/- 30.34% and 49.64 +/- 25.74% at 5 m, 1 h and 24 h, respectively. MPI or BLI could not identify the bacteria in the tumor in live animals. However, after excision of the tumor, iron was detected *ex vivo* in 2/5 tumors using MPI (**Figure 5b**). One tumor was cut in half immediately after excision and luminescent signals were identified in the tumor core (**Figure 5c**); signals were not identified in intact tumors.

**Figure 5.**
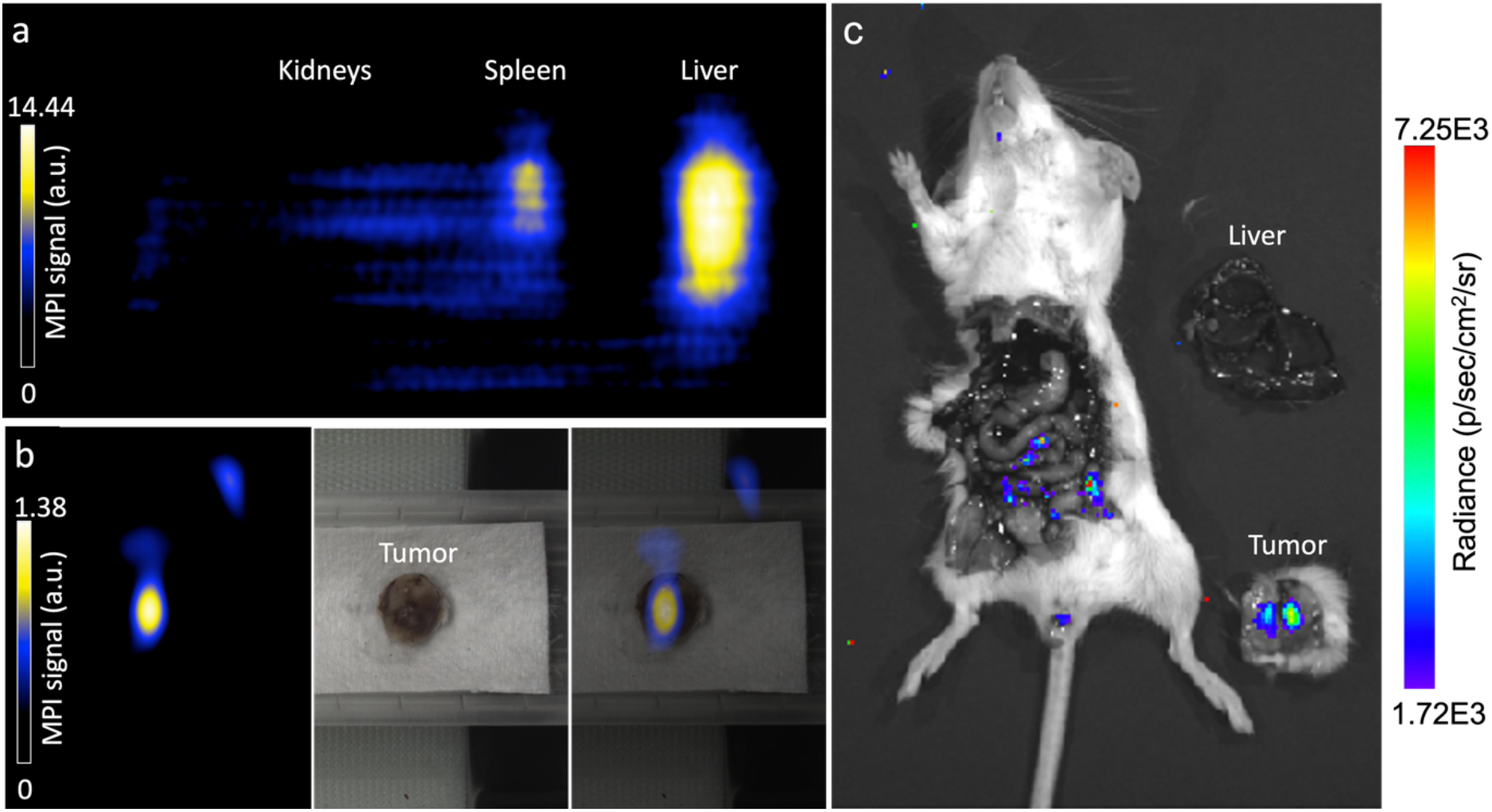
Excised organs and postmortem MPI and BLI. Ex vivo MPI of the kidneys, spleen and liver identifies signal in the liver and to a lesser extent the spleen (a; healthy mouse). MPI of an excised tumor identifies MPI signal (b). IVIS imaging of a mouse postmortem and excised liver and tumor identify signals in the intestine and within the core of the tumor (c).

In this study, we monitored viability of a bacterial contrast agent *in vivo* by imaging bioluminescent signals and used these images to guide analysis of iron accumulation within tissues before and after the loss of bacterial viability. Revealing the temporal localization and integrity of this living magnetic contrast agent will enable its development as an imaging agent and a delivery vehicle as a bacteriotherapy. The bioluminescent signals dispersed throughout the mouse immediately after IV injection indicated successful systemic delivery. The decrease in luminescence could be due to both bacterial death as well as poor penetration of the 490 nm bluegreen light produced by *luxA-E*, as was noted when signals were detectable when an excised tumor was cut in half. However, the magnetosomes in the magnetotactic bacteria can still be used as an MPI contrast agent, providing information on location and concentration even after the bacteria are killed. Combining BLI with MPI can provide anatomic localization of viable bacteria which can be correlated to the localization of magnetosomes.

Magnetotactic bacteria may be better as drug delivery systems, versus passive agents such as extracted magnetosomes or nanoparticles, following IV or peritumoral injection, presumably due to viable agents reaching deeper into the tumor^38^. It is equally plausible that the relatively few bacteria that reach the tumor can replicate in the immune suppressive environment of the tumor bed. It may be this amplification through replication that enables persistence and penetration into the tumor^46–48^. The Gram-negative bacterial lipopolysaccharide (LPS), which is produced by MSR-1, is known to produce an immunological response^49^; although this may facilitate immune rejection of the tumor, it may also lead to the systemic elimination of MSR-1 thus reducing both background signals in imaging and off-target effects in therapy.

### Fluorescence microscopy of excised tumor and liver

Fluorescence microscopy identified Lipid A LPS (green), potentially as intact or pieces of Gram-negative bacteria, within the tumor and liver sections (**Figure 6**). These regions were associated with both F4/80+ (red; macrophages, closed arrowhead) and F4/80-cells (open arrowhead, **Figure 6a**). Lipid A LPS structures were identified in both the tumor (**Figure 6b**) and liver (**Figure 6c**) appearing as intact elongated structures, and also identified in the tumor, not associated with any cell (**Figure 6d**).

**Figure 6.**
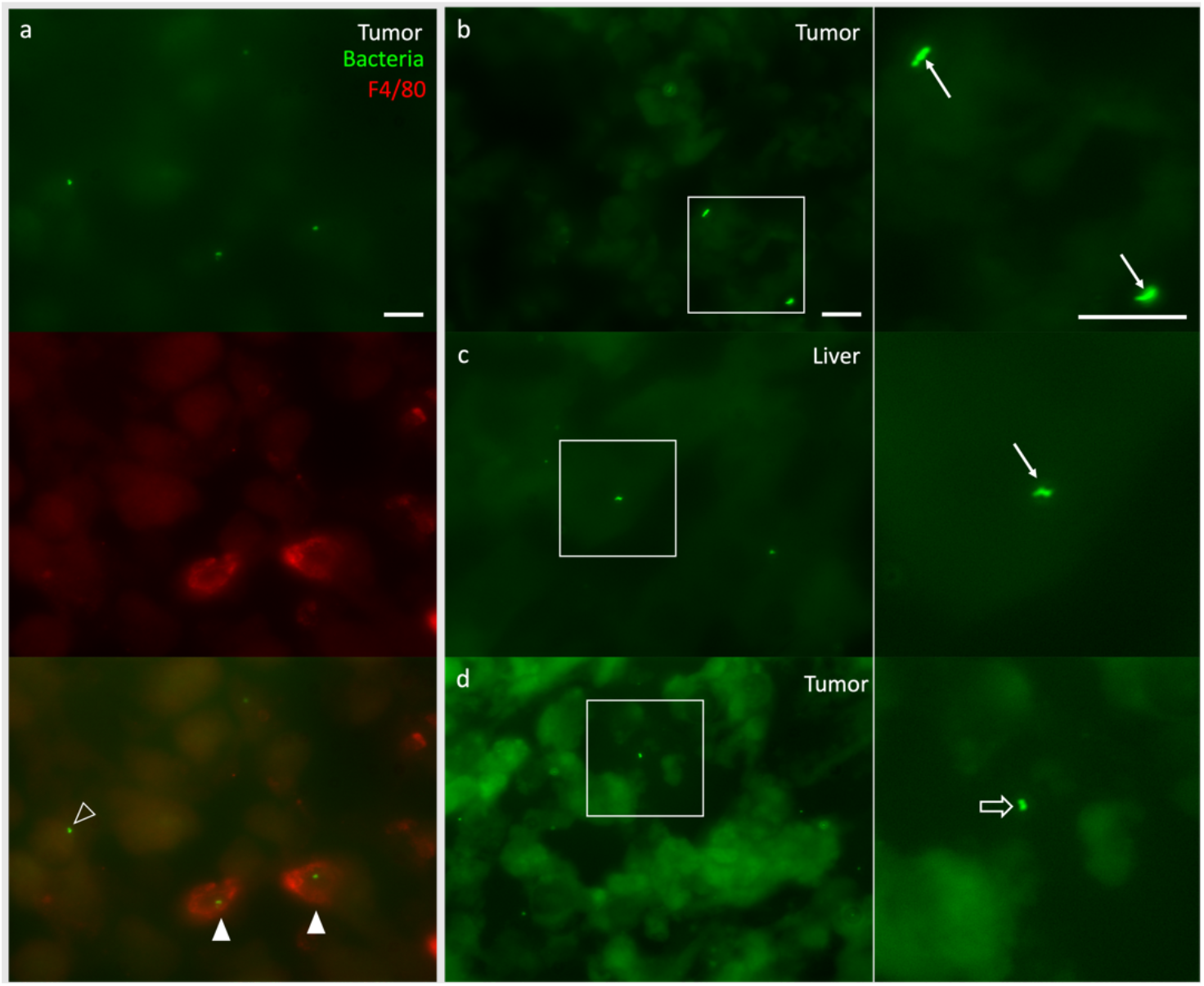
Histological analysis of excised tumor and liver. Within the tumor, bacteria (green) are identified in the same region as F4/80 positive (red) cells (solid arrowheads), as well as F4/80 negative cells (open arrowhead, a). Within the tumor (b) and liver (c), bacteria are identified which appear to have retained their morphology (white arrows, zoomed insets adjacent). Within the tumor there were bacteria which were not associated with a cell (d, open arrow, zoomed inset adjacent). Scale bars = 20 *μ*m.

The Lipid A LPS+ antibody could identify other Gram-negative bacteria within the tumor and therefore it cannot be certain that the bacteria identified are the MSR-1 which were injected. However, the presence of iron by MPI, and bioluminescence using BLI ensures that there are bacteria in these locations. Many groups have demonstrated tumor-targeting bacteria *in vivo*. Although a systemic injection results in distribution to normal tissues as well as the tumor, bacteria in regions unrelated to neoplasia will clear within hours to days, while those in the tumor continue to be viable and even proliferate at the tumor site^46–48,50^. This is likely due to the immunosuppressive environment of the tumor, and possibly necrotic and hypoxic tumor cores, as well as the unique biochemical environment^51,52^. These studies have used pathogenic bacteria including *Salmonella spp., Listeria spp*. and *Clostridium spp*., and when investigating an anti-tumoral response, rely on extracellular interactions. The response of magnetotactic bacteria in the tumor is less well understood. However, *Magnetococcus marinus* MC-1 was shown to deliver therapeutics to tumors^38^ and the addition of improved MPI tracking coupled with bioluminescence would provide enhanced understanding accumulation and longevity *in vivo*. The loss of viability seen with MSR-1 *in vivo* appeared to be more rapid than that observed for MC-1, and therefore selection of the delivery organisms can be guided by multimodality imaging.

If we were to identify magnetotactic bacterial persistence within the tumor microenvironment, we would be presented with a unique opportunity to use bacteriotherapy to target cancer. The imaging properties would allow for visualization of localization and persistence, and the bacteria could produce therapeutic effects. For example, magnetic hyperthermia could be used to ablate cells by heating the magnetosomes or further, the magnetosomes could be heated to a lesser extent, thermally inducing the expression a gene of interest to elicit anti-tumoral effects.

## Conclusion

The results provided here offer a link between structure of nanoparticles, microbiology and imaging modalities. The significance of this study comes from comparing the mutant and wildtype forms of these biological nanoparticles via MPI and the conclusion that clustered aggregates of biological nanoparticles resulted in improved signal. Exploring how the physical rotations of these biological nanoparticles vary because of shape or arrangement of these magnetosomes/particles would be useful to further understand the physics of iron nanoparticle imaging. Our results describe the application of magnetotactic bacteria for MPI and creates many possibilities of future use as living contrast for *in vivo* imaging and for other synthetic biology applications. There is potential for using magnetotactic bacteria for a variety of applications including bacteriotherapy, drug delivery, hyperthermia and as contrast agents, and improving functional imaging with MPI will enable advancement of these applications.

## Methods

### Cultivation and Growth of MSR-1

*M. gryphiswaldense*, strain MSR-1 (wild-type: MSR-1 WT; and MamJ deficient mutant: MSR-1 Δ*mamJ* (Dirk Schüler, University of Bayreuth))^27^ were grown in an environmental-controlled chamber at 1% O_2_, 5% CO_2_ and 30°C on a rotary shaker at 230 rpm. Cells were cultured in growth medium containing 10 mM HEPES pH 7.0, 4 mM NaNO_3_, 0.6 mM MgSO_4_ · 7H_2_O, 0.74 mM KH_2_PO_4_, 3 g/L Soybean peptone, 0.1 g/L Yeast Extract and 15 mM Potassium L-Lactate. After sterilization, the iron source Fe(III) citrate was added at a concentration of 50 *μ*M. Selection of the *luxA-E*^33^, expressing strains were maintained using media supplemented with 15 *μ*g/mL kanamycin. Optical density of cultures was determined photometrically at 565 nm.

### Transmission Electron Microscopy

Transmission Electron Microscopy (TEM; JEM-1400Flash, JEOL, MA USA) was used to confirm morphological differences between the Δ*mamJ* and WT strains. Samples were fixed in 2.5% EM-grade glutaraldehyde for 5 m. Ten microliters (*μ*l) of fixed sample were incubated for 10m on 200-mesh, carbon-coated copper grids. Following incubation, the grids were washed three times with HPLC-grade water and imaged. Images were analyzed using Fiji (ImageJ version 2.0.0-rc-69/1.52i).

### luxA-E Constructs

*Escherichia coli* mating strain WM3064 was grown in Luria Broth with 0.3 mM Diaminopimelic acid (DAP) at 37°C. *E. coli* WM3064 containing the pAK532 mobilization plasmid was kindly provided by Arash Komeili (UC Berkeley)^25^. The *luxA-E* operon was cloned into the pAK532 linearized backbone using Gibson Assembly^42^ and transformed into the *E. coli* WM3064 mating strain (Supplemental Table 1). Colonies constitutively expressing the enzymes for bioluminescence were selected on 50 *μ*g/ml kanamycin and confirmed by restriction enzyme digests. The *E. coli* WM3064 pAK532 *luxA-E* strain was conjugated with MSR-1 WT and Δ*mamJ* to create a constitutively bioluminescent strain of MSR-1 (MSR-1 WT *luxA-E* and MSR-1 Δ*mamJ luxA-E*, respectively). Two hundred microliters of WM3064 and MSR-1 Δ*mamJ* or MSR-1 WT were mixed, centrifuged and resuspended in 10 ml and added to plates containing agar-solidified media + 0.3 mM DAP. Plates remained at room temperature under 7% O_2_ for 1 h and then moved to an anaerobic chamber with 1% O_2_ and 5% CO_2_ for 4 h. The mixture was then spread on the surface of a plate with kanamycin selection (15 *μ*g/ml). Plates were placed in an anerobic jar with palladium-coated pellets at 30°C and grown for 5 days, when bioluminescent colonies were selected.

### Inductively Coupled Plasma-Mass Spectrometry (ICP–MS)

Δ*mamJ luxA-E* and WT *luxA-E* cells were washed with a 20 mM Hepes-4 mM EDTA solution and 8.0×10^8^ cells (n = 3) were collected. Following MPI (below), the cells were digested in concentrated nitric acid (J.T. Baker, USA; 69-70%) overnight, and diluted 25-fold with a solution containing 0.5% EDTA and Triton X-100, 1% ammonium hydroxide, 2% butanol, 5 ppb of scandium, and 7.5 ppb of rhodium, indium, and bismuth as internal standards (Inorganic Ventures, VA, USA). The samples were analyzed on an Agilent 7900 ICP mass spectrometer (Agilent, CA, USA). Elemental concentrations were calibrated using a 5-point linear curve of the analyte-internal standard response ratio. Bovine liver (National Institute of Standards and Technology, MD, USA) was used as a control.

### In Vitro Imaging

MPI (Momentum, Magnetic Insight Inc, CA, USA) was acquired using a 2D projection scan with default settings (5.7 T/m gradient, 20 mT, 45 kHz excitation) and the following imaging parameters: field of view (FOV) = 4 x 6 cm, 1 average and acquisition time of 15 seconds (s). MPI was performed on the same Δ*mamJ* or WT *luxA-E* pellets used for ICP-MS (prepared as described above) and iron content was compared to ICP-MS values.

BLI (IVIS, PerkinElmer; exposure time = 2-3 s, binning = medium, f/stop = 1, emission filter = open) was performed on 2.5×10^8^, 1×10^8^ and 0.5×10^8^ cells of Δ*mamJ* or WT *luxA-E* (n = 3), followed by MPI (as above, with 2 signal averages). MPI and bioluminescent signals were compared, between strains and number of cells.

Relaxometry was performed on 2.85×10^9^ Δ*mamJ* and WT *luxA-E* pellets. Signal amplitude (peak signal strength) and spatial resolution (calculated using full width at half maximum, FWHM) were calculated using the information from the PSF generated.

### Tumor Cells and Animal Model

Female BALB/c mice (6-8 weeks; Charles River USA) were obtained and cared for in accordance with the standards of Michigan State University Institutional Animal Care and Use Committee. 4T1 murine mammary carcinoma cells (American Type Culture Collection, USA) were maintained in RPMI containing 10% fetal bovine serum and 50 U/ml Penicillin and 50 *μ*g/ml Streptomycin at 37°C and 5% CO_2_. Mice (n = 5) were anesthetized with isoflurane administered at 2% in oxygen followed by an injection of 300,000 4T1 cells (>90% viability, measured using the trypan blue exclusion assay) in 50 *μ*l phosphate buffered saline (PBS). Injections were into the 4^th^ (inguinal) mammary fat pad (MFP), as previously reported^43^.

### In Vivo Imaging

BLI, MPI and CT imaging were performed at 5 m, 1 h and 24 h post IV injection of 2.1×10^9^ cells of Δ*mamJ luxA-E* bacteria in PBS into healthy (n = 3) or 4T1 tumor-bearing (n = 5) mice. First, BLI was performed using an IVIS Spectrum system (PerkinElmer; exposure time = auto (up to 300 s), binning = medium, f/stop = 1, emission filter = open). The mouse was then transferred into the MPI bed where iron fiducial markers were placed in 3 dimensions to allow for co-registration with CT images. 2D and 3D MPI tomographic scans were performed using the following parameters: FOV: 12 x 6 x 6 cm, 5.7 T/m gradient, 1 (2D) or 21 (3D) projections and 1 average. The bed was then transferred to a Quantum GX microCT scanner (PerkinElmer). Whole body CT images were acquired using 3 x 8 s scans with the following parameters: 90 kV voltage, 88 *μ*A amperage, 72 mm acquisition FOV and 60 mm reconstruction FOV resulting in 240 *μ*m voxels. After the final imaging time point mice were sacrificed and organs of interest were excised and imaged by BLI and MPI followed by immunofluorescence staining and microscopy (below).

### Image Analysis

MPI data sets were visualized and analyzed utilizing Horos imaging software. To calculate total MPI signal for quantification purposes, a fixed-size region of interest (ROI) was used for all data sets (pellets, *in vivo*, *ex vivo* organs). The size of ROI was chosen to accommodate the largest signal area (liver signal). Any regions of negative signal were set to zero. Total MPI signal was calculated by *mean signal × volume* and displayed in arbitrary units (a.u.).

BLI data were analyzed using Living Image software (PerkinElmer, Version 4.5.2). ROI were placed around tube or mouse and average radiance of the pixels in the ROI is reported as radiance (p/sec/cm^2^/sr).

### Histological Analysis

After imaging of excised organs (liver, spleen, lymph nodes) and tumors, samples were fixed in 4% paraformaldehyde for 24 h followed by cryopreservation through serial submersion in graded sucrose solutions (10%, 20% and 30%). Sections were then frozen in optimal cutting temperature compound (Fisher HealthCare, USA). Tissues were sectioned using a cryostat (10 *μ*m sections). Sections were then stained with primary F4/80 Monoclonal Antibody (BM8), (Thermo Fisher Scientific, catalog #14-4801-85) and Lipid A (Novus Biologicals, CO USA, catalog #64484) and secondary Goat anti-Rat IgG (H+L), Alexa Fluor 647 (Thermo Fisher Scientific, catalog #A-21247) and Donkey anti-goat IgG (H+L) Alexa Fluor 488 (AbCam, catalog #ab150129). Tissue sections were imaged using a Leica DMi8 Thunder microscope equipped with a DFC9000 GTC sCMOS camera and LAS-X software (Leica, Wetzlar, Germany). Fluorescence images were prepared using Fiji (ImageJ).

### Statistical Analysis

Statistical analyses were performed using Prism software (9.0.0, GraphPad Inc., CA, USA). Iron content/cell and MPI signal/*μ*g iron data were analyzed using an unpaired t-test. Linear regression was performed on MPI and BLI data performed on cells for correlation. Percent of injected dose for BLI and MPI at each time point were analyzed using a repeated measures one-way ANOVA followed by Tukey’s multiple comparisons test. Data are expressed as mean +/- standard deviation; *p*<0.05 was considered a significant finding.

## ASSOCIATED CONTENT

### Supporting Information

The following files are available free of charge.

Supplemental Table 1 – List of oligonucleotides used in this study.

## AUTHOR INFORMATION

### Author Contributions

The manuscript was written through contributions of all authors. All authors have given approval to the final version of the manuscript.

## ACKNOWLEDGMENTS

The Contag Lab would like to acknowledge the Schüler lab for providing the original MSR-1 Δ*mamJ* mutant strain, and the Komeili lab at the University of California Berkeley for the gift of pAK532 and *E. coli* WM3064 as well as various magnetotactic bacteria protocols and advice. We would also like to acknowledge the MSU Veterinary Diagnostic Lab for ICP-MS and Dr. Alicia Withrow at the MSU Center for Advanced Microscopy for TEM protocols and imaging. The authors would like to thank the James and Kathleen Cornelius Endowment for funding of this project. AVM was funded by a Natural Science and Engineering Research Council Postdoctoral fellowship for this project.

