## Supplemental Table 1 for "Magnetic particle imaging of magnetotactic bacteria as living contrast agents is improved by altering magnetosome structures"

| Primer Name | Sequence | |
| --- | --- | --- |
| Lux_pAK532_primer | FWD | AATTTCACACAGGAAACAGAATTCATGAAGCAAGAGGAGGACTCTCTATG |
|  | RVS | GCCGCTCTAGAACTAGTTCATTAACTATCAAACGCTTCGGTTA |
| pAK532_linearization_primer | FWD | TGAACTAGTTCTAGAGCGGC |
|  | RVS | GAATTCTGTTTCCTGTGTGAAATT |

**Supplemental Table 1.** List of oligonucleotides used in this study.
